# Differentially polarized macrophages show diverse proangiogenic characteristics under normo- and hyperglycemic conditions

**DOI:** 10.1101/2024.01.12.575474

**Authors:** Mahnaz Shariatzadeh, César Payán Gómez, Willem A. Dik, Pieter J. M. Leenen

## Abstract

**Purpose:** Angiogenesis is a vital process involved in the formation of new blood vessels from existing ones. Macrophages play a crucial role in initiating endothelial activation and inflammation, and are involved in the pathological angiogenesis. Traditionally, macrophages have been classified, with the pro-angiogenic activity attributed to the M2 phenotype. However, recent evidence challenges the notion that only M2 macrophages possess pro-angiogenic properties. This study aims to investigate the pro- and anti-angiogenic properties of human polarized macrophages in normo- and hyper-glycemic conditions, in order to gain a better insight into the angiogenic capacity of M1- and M2-like macrophages in diabetes.

**Methods:** A comprehensive bioinformatic analysis of pro- and anti-angiogenic gene expression profiles related in M1-vs. M2-polarized macrophages was performed based on a large previously published dataset. The most contributing differentially expressed genes in angiogenesis were selected for further validation. Macrophages were generated and polarized by culturing CD14^+^ monocytes and their stimulation with any of IFN-γ, IL-4, or IL-6 cytokines. Polarized macrophages were immunophenotyped using flow cytometry, and their expression of the selected genes were measured using qPCR. Finally, the proangiogenic capacity of the cells was assessed in an *in vitro* 3-D endothelial tubule formation assay, containing GFP-expressing human retinal endothelial cells, pericytes, and pro-angiogenic growth factors.

**Results:** IL-4 and IL-6 induce distinct M2-like phenotypes in macrophages with mixed pro- and anti-angiogenic gene expressions. Hyperglycemia has a mild negative effect on the expression of M2-associated markers, however it does not significantly affect the angiogenic properties of macrophages.

**Conclusion:** Our data support the concept of a spectrum model for macrophage polarization, indicating that the angiogenic status of polarized macrophages is not limited to the M2-phenotype, but is rather mediated by microenvironmental cues, and can result in diverse phenotypic characteristics. The effect of hyperglycemia on the angiogenic capacity of macrophages requires more comprehensive investigation.

## Introduction

Angiogenesis is the crucial physiological process through which new blood vessels are formed from pre-existing vasculature ^1^. Under normal physiological conditions, angiogenesis is tightly controlled and requires strict balance between the pro-angiogenic and anti-angiogenic factors^2^. However, under certain circumstances when pro-angiogenic activity predominates over anti-angiogenic activity, pathological angiogenesis occurs, as for instance seen in different types of cancer, rheumatoid arthiritis and diabetic retinopathy (DR). There is increasing evidence that pathological angiogenesis, characterized by abnormal blood vessel formation, is associated with inflammation ^1^. Monocytes/macrophages play a pivotal role in triggering endothelial activation and inflammation, for instance in cardiovascular disease and progression of atherosclerosis ^3^. Additionally, monocytes/macrophages contribute to pathological angiogenesis observed in tumors.

The diversity in the function of macrophages is ascribed to the plasticity and ability to differentiate into distinct subsets with varying properties ^4^. Based on the micro-environmental cues, myelomonocytic cells undergo polarization into a flexible spectrum of different polarities, with tissue- and context-specific properties ^5^. According to the traditional nomenclatures, *in vitro* classical macrophage activation corresponds to *in vivo* M1-like phenotype, while alternative activation is associated with M2-like macrophages. Several studies have been conducted to identify cytokines/chemokines and cytokines that are more specifically associated with either M1 or M2 macrophages ^6, 7^. These studies have linked pro-inflammatory molecules (eg. IL-1β, IL-6, and TNF-α) and pathways to the M1 phenotype, while anti-inflammatory factors (eg. IL-10, IL-13, and TGF-β) have been associated with M2-like macrophages. However, it has been noted that additional subsets of macrophages exist that appear as intermediates between M1 and M2 macrophages and express both M1- and M2-related genes ^8^. This indicates that the conventional model of M1 and M2 polarization may be oversimplified, and that the “classical” M1 and “non-classical” M2 macrophage subtypes likely represent the extremes of a spectrum ranging from pro-inflammatory to an anti-inflammatory state. Moreover, the paradigms of macrophage M1(classical) vs. M2(alternative) activation is no longer accepted by new research as there is a small overlap between the gene expression profiles of *in vivo* M1/M2 and *in vitro* classically/alternatively activated macrophages ^4, 5^.

The conventional classification also assigns the pro-angiogenic activity of macrophages to M2 phenotype, which is traditionally assumed to attenuate inflammation and participate in tumor progression and tissue remodeling ^9, 10^. This alignment with the understanding that tissue remodeling and trophic capacities are typically associated with M2-like polarization seems logical at first glance. However, upon closer examination, we question the generalization of this notion and suggest a more nuanced perspective. Emerging evidence challenges the exclusive association of pro-angiogenic activity with the M2 phenotype ^11^. It has been observed that the pro-inflammatory state linked to M1-like polarization of macrophages plays a role in pathological angiogenesis, specifically known as inflammatory angiogenesis ^11^. This finding expands the range of macrophage subsets involved in angiogenesis beyond the M2 phenotype. Moreover, recent research conducted in mice has demonstrated that retinal macrophages and microglia involved in pathological angiogenesis exhibit a highly glycolytic metabolism ^12^, typically associated with M1-like macrophages ^13^, although they show features of both M1 and M2 polarization. This suggests that macrophages engaged in angiogenesis can possess mixed phenotypes and metabolic profiles, challenging a clear-cut classification.

In the context of diabetes, there is evidence of a low-grade chronic inflammation caused by the recruitment and accumulation of monocytes and granulocytes in to the inflamed tissues (such as retina), which has been associated with vascular dysfunction ^14^. Hyperglycemia, a hallmark of diabetes, induces a mixed phenotype differentiation of macrophages into both M1- and M2-like subpopulations ^15^. Both of these subpopulations contribute to the development of chronic inflammation in endothelial cells, leading to the occurrence of macrovascular complications such as atherosclerosis, as well as microvascular complications such as DR.

In this study, we aim to identify the pro- and anti-angiogenic factors expressed during human macrophage activation and polarization in normo- and hyper-glycemic conditions, in order to gain a better insight into the angiogenic capacity of M1- and M2-like macrophages in diabetes. To achieve this, we performed a comprehensive bioinformatic analysis of the gene expression profiles related to pro- and anti-angiogenic genes in M1-vs. M2-polarized macrophages, based on a large publically available dataset ^15^. To substantiate our findings, we have differentiated and polarized macrophages into M0-, M1-, and M2-like macrophages under normal or high glucose conditions, and conducted an unbiased validation of the differentially expressed genes (DEG) retrieved from aforementioned data sets that had the highest contribution in promoting or inhibiting angiogenesis. Furthermore, we evaluated the proangiogenic capacity of the polarized macrophage subsets by examining their effects on *in vitro* tubule formation of retinal endothelial cells (REC), with the aim of replicating the pathological angiogenesis observed in DR.

## Materials and methods

### Recompilation and analysis of macrophage datasets

A systematic search was performed in the Gene Expression Omnibus Database (https://www.ncbi.nlm.nih.gov/geo/) to identify studies that have analyzed the genome-wide expression of human macrophages exhibiting M0, M1, and M2 phenotypes in the context of hyperglycemia. The experimental protocol required the adherence to the following criteria: isolation of monocytes from buffy coat, induction of differentiation towards M1 and M2 phenotypes, and subsequent cultivation in the presence of normal and high glucose concentrations. The transcriptomes of the cells were measured using the Affymetrix Human Gene 1.0 ST Array ^15^, resulting in 24 arrays with four biological replicates for M0, M1, and M2 cultivated under both-NG and-HG conditions. In our analysis, all arrays passed the quality control assessments, exhibiting similar RNA degradation plots and intensity distributions prior to normalization, with no detected outliers in the PCA using raw data.

The transcriptomic data analysis was conducted through a series of steps. First, all samples underwent quality control using the QC module from simpleaffy ^16^, which utilized various parameters such as virtual image reconstruction, signal comparability, and array correlation to identify low-quality samples. Data preprocessing was performed using the Limma R/Bioconductor software package ^17^, wherein probesets were summarized, data were normalized, and log2 transformed using the robust multichip average (RMA) algorithm.

To identify differentially expressed genes (DEG), the linear model from Limma ^17^ implemented in R was used. Pairwise comparisons were made between M1 vs M0, and M2 vs M0 samples, and the fold change (FC), *P* value, and false discovery rate (FDR) were calculated for each probe in the microarrays. A cut-off value for DEG was set at FDR <0.05 with a fold change ≥ |1.5|.

### Collection of pro- and anti-angiogenic signatures

We generated a comprehensive list of genes (Suppl. Table 1) that were annotated with either pro-angiogenic or anti-angiogenic activity. Initially, we conducted a search for genes that were annotated in either of these two categories, utilizing both Ingenuity Pathway Analysis (IPA)^18^ and the Gene Ontology Biological Processes databases.

### Principal component analysis of pro- and anti-angiogenic genes in M0, M1 and M2 macrophages

Principal component analysis (PCA) was computed to identify whether genes annotated in anti- and pro-angiogenic signatures exhibited distinct patterns of expression among the M0, M1, and M2 samples. The following steps were undertaken for each signature (anti-angiogenic and pro-angiogenic): first, the expression levels of the various genes within each signature were selected for all the samples; second, PCA was computed; third, genes that contributed significantly to the separation of M0, M1, and M2 samples in the PCA plot were identified; and finally, the DEG were extracted from the list of genes obtained in the previous step.

To identify the most influential genes in the clustering of the M1, M2, and M0 samples, we used the pcaGoPromoter package implemented in R ^19^. We calculated the loads (genes) for each principal component, allowing for the ranking of the genes based on their importance in the distribution of the samples across the principal components.

### PBMC isolation

Peripheral blood mononuclear cells (PBMC) were isolated from normal blood donor buffy coats using standard two-step Ficoll-Paque gradient centrifugation ^20^. Cell pellets were resuspended in RPMI freezing medium (containing 40% fetal bovine serum (FBS, Gibco) + 10% Dimethyl sulfoxide (DMSO, Sigma-Aldrich, St Louis, MO, USA)) and stored in liquid nitrogen until further use.

### Human monocyte isolation

PBMC were thawed, counted (Countess II; Thermofisher,Waltham,MA, USA), and stained with MACS colloidal superparamagnetic microbeads (# 130-050-201, Miltenyi Biotec, Leiden, the Netherlands) conjugated with monoclonal anti-human CD14 antibodies for positive selection of CD14^+^ monocytes according to manufacturer’s instructions. The purity of CD14^+^ cells (> 95% pure) was confirmed using flow cytometry.

### Macrophage polarization

Monocytes were cultured in RPMI 1640 medium (Gibco) supplemented with 10% FBS (Gibco) on 12-well (1 × 10^6^ cells/ml) and 48-well (0.4 × 10^6^ cells/200 µl) tissue culture plates for functional assays and flowcytometric analysis, respectively. From day zero, the cells were stimulated with either M-CSF (100 ng/ml, # 130-093-491, Miltenyi Biotec), M-CSF + IL-4 (40 ng/ml, # 130-093-917, Miltenyi Biotec), M-CSF + IL-6 (50 ng/ml, # 206-IL-050/CF, R&D systems), or M-CSF + IFN-γ (40 ng/ml, # 285-IF-100/CF, Bio-Techne) in normal (11 mM), or high (25 mM) glucose concentrations. After 4 days of culture, the morphology was examined at 20× magnification using an Axiovert 100 light microscope (Zeiss, Oberkochen, Germany). Next, the culture supernatant was discarded and adhered cells were harvested for flow cytometric analysis using 30 minutes incubation with cold 1 mM PBS (calcium- and magnesium-free) + EDTA, or prepared for mRNA isolation.

### Flow cytometry

The harvested cells were washed with staining buffer (MACSima^TM^ running buffer, Miltenyi Biotec, Bergisch Gladbach, Germany) and exposed to anti-human monoclonal fluorescent antibodies at 4⁰C in a dark environment according to the manufacturer’s instructions. The following antibodies were used: CD14-PE-Cy7 (61D3, eBioscience^TM^), CD16 (FcγRI)-APC-Cy7 (3G8 (RUO), BD Pharmingen^TM^), CD163-PE (GHI/61, BD Pharmingen^TM^), CD64-APC (10.1 (RUO), BD Pharmingen^TM^), anti-HLA-DR (MHC class II)-FITC (G46-6, BD Pharmingen^TM^) and CD206 (MRC1)-FITC (15-2; BioLegend). After 20 minutes of incubation, the cells were washed and resuspended in 200 µl staining buffer. Surface marker expression was measured using FACScanto^TM^II cell analyzer (BD Biosciences, Piscataway, NJ, USA). Viable cells were gated and data were analyzed using FlowJo software (Tree Star, Ashland, OR, USA). At least 10,000 events were acquired per sample.

### Real-time quantitative polymerase chain reaction analysis

After 4 days of macrophage differentiation, mRNA was isolated and converted into cDNA using a commercially available kit (GenElute Mammalian Total RNA Miniprep Kit; Sigma-Aldrich, USA). Transcript levels were determined by real-time quantitative polymerase chain reaction (QuantStudio 5; Thermo Fisher Scientific, Waltham, MA), normalized to the control gene Abelson (*ABL*) and expressed as relative mRNA expression using the ΔCT method ^21^. Primer-probe combinations used are listed in Suppl. Table 2.

### 3-D *in vitro* tubule formation assay

To investigate the impact of polarized macrophages on REC vessel formation, we utilized a three-dimensional *in vitro* tubule formation assay as described previously ^22, 23^. In this assay, primary human retinal endothelial cells (REC: cat # ACBRI 181, Cell Systems, Troisdorf, Germany) transduced with GFP-tagged lentiviral vector were co-cultured with human brain vascular pericytes (cat # SCC1200-KIT, ScienCell research, San Diego, USA) in a collagen matrix (bovine collagen type-1, Gibco^TM^, cat # A1064401) in the presence of three pro-angiogenic growth factors (IL-3, stem cell factor (SCF), and stromal cell-derived factor 1-alpha (SDF1-alpha); R&D systems, Abingdon, UK; 25–200 ng/ml as indicated). After 24 hours, successful sprouting of endothelial cells was typically observed, and the differentiated macrophages were added on top of the wells in four replicates (500 cells/well). After 4 days of co-culture, tubule formation of the REC was visualized using ×20 magnification of an inverted fluorescence microscope (Olympus SC30, Shinjuku, Japan), and the total surface area was quantified using FIJI software (version 1.51n).

### Statistics

Data obtained from flow cytometry, qPCR, and tubule formation assay were analyzed using GraphPad Prism (version 5.04). To determine significant differences between different experimental conditions, student T-test, non-parametric Mann Whitney U T-test or one-way ANOVA followed by a post hoc Tukey multiple comparison test were used, and a *P* value < 0.05 was considered statistically significant. All data are presented in means ± standard error of the mean (SEM).

## Results

Our search in the GEO omnibus database found one study that investigated human macrophage polarization in the context of hyperglycemia, and thus met our criteria: GSE86298. This dataset was derived from monocytes from four healthy human donors, isolated using CD14 beads from buffy coats, which were then stimulated for 6 days with IFN-g (M1), IL-4 (M2), or no additional cytokines (M0). These cells were cultivated in the presence of either 5 mM (normal glycemia, NG) or 25 mM (hyperglycemia, HG) glucose. The transcriptomes of the cells formed the basis of a publication describing the influence of hyperglycemia on parameters of polarization ^15^.

Given our interest in examining the response of pro- and anti-angiogenic genes in M1- and M2-like macrophages and the impact of glucose concentration on the transcriptome, we utilized the data on M0 macrophages cultivated under-NG conditions as a reference. To identify if angiogenesis-related genes were modulated in the process of differentiation of M1 or M2 from M0 we calculated the lists of differentially expressed genes (DEG) between M1-NG vs. M0-NG, and M2-NG vs. M0-NG. We subsequently computed the lists of DEG when comparing M1 and M2 under NG and HG conditions (Suppl. Table 3).

Based on the data presented in Suppl. Table 3, the number of DEG and the ratio of up-regulated and down-regulated genes observed in M1 and M2 after stimulation from M0 were comparable. However, exposure to high levels of glucose resulted in partial modifications to the transcriptome. Specifically, in M1, 563 genes (1753 minus 1190; 32% of DEG) were altered in M1-HG vs M0-NG, and in M2, 233 genes (1421 minus 1188; 26% of DEG) were DEG in M2-HG vs M0-NG but not in M2-NG vs M0-NG.

### Identification of the of pro- and anti-angiogenic genes regulated upon M1 and M2 macrophage polarization

The biological interpretation of the lists of DEG was performed using the over-representation analysis implemented in DAVID, with specific focus on the involvement of angiogenesis-related pathways. Firstly, we analyzed the list of DEG induced by M1 differentiation under NG and HG conditions (Suppl. Table 4A and 4B, resp.). Interestingly, ‘Angiogenesis’ (GO:0001525) was the eighth most prominent pathway with 38 DEG and an adjusted *P* value equal to 0.0024 (Suppl. Table 4A). Another angiogenesis-related pathway, with a statistically significant *P* value was the ‘Positive regulation of angiogenesis’ (GO:0045766) with 22 DEG and an adjusted *P* value of 0.0262.

It appeared that pathway analysis of the DEG for M1 under HG conditions, when compared to M0, yielded results similar to those found under NG conditions, with a predominance of regulated immune system pathways. However, compared to M1-NG, the angiogenesis pathway showed a smaller adjusted *P* value (4.05E-04 vs. 0.0024), associated with a higher number of DEG (41 vs. 38), and was identified as second most important among the identified pathways (Suppl. Table 3B).

Pathway analysis of DEG for M2 in NG and HG conditions compared to M0 showed similar results, with the immune system being the most prominent pathway affected (Suppl. Table 5). However, some differences with the M1 results were apparent, as the angiogenesis pathway was ranked lower in the M2 top 10 of affected pathways. Additionally, there were no significant differences in the position, number of genes, and adjusted *P* value for angiogenesis in M2 cultivated with NG or HG (Supplementary Table 5). Pathway analysis indicated that angiogenesis-related genes are modulated upon both M1 and M2 polarization of macrophages in a quantitatively similar fashion. This occurs comparably under normal or elevated glucose levels, albeit the hyperglycemic effect on angiogenic activity might be somewhat stronger upon M1 polarization.

Next, we investigated whether the expression levels of genes annotated as pro- or anti-angiogenic were able to distinguish cells based on their polarization state, and whether glucose levels influenced this. Therefore, we performed PCA analysis, using a selected set of pro- and anti-angiogenic genes, as annotated in the Gene Ontology and Ingenuity Pathway Analysis databases. The pro-angiogenic pathway consisted of 302 genes, while the anti-angiogenic pathway had 114 genes. Figure 1 shows the PCA result for cellular expression profiles using pro-angiogenic (Fig. 1A) and anti-angiogenic genes (Fig. 1B), as determined under normoglycemic conditions. Concerning pro-angiogenic genes, in PC1 (representing 65% of diversity), M0 cells are intermediate between M1 and M2 cells. With regard to anti-angiogenic genes, however, M0 and M2 cells show a similar PC1 position (80% of diversity), while M1 cells stand apart in this respect. These results indicate that M1 and M2 cells exhibit clear differences in the levels of angiogenesis-related gene expression compared to each other and to M0, as expected.

**Figure 1.**
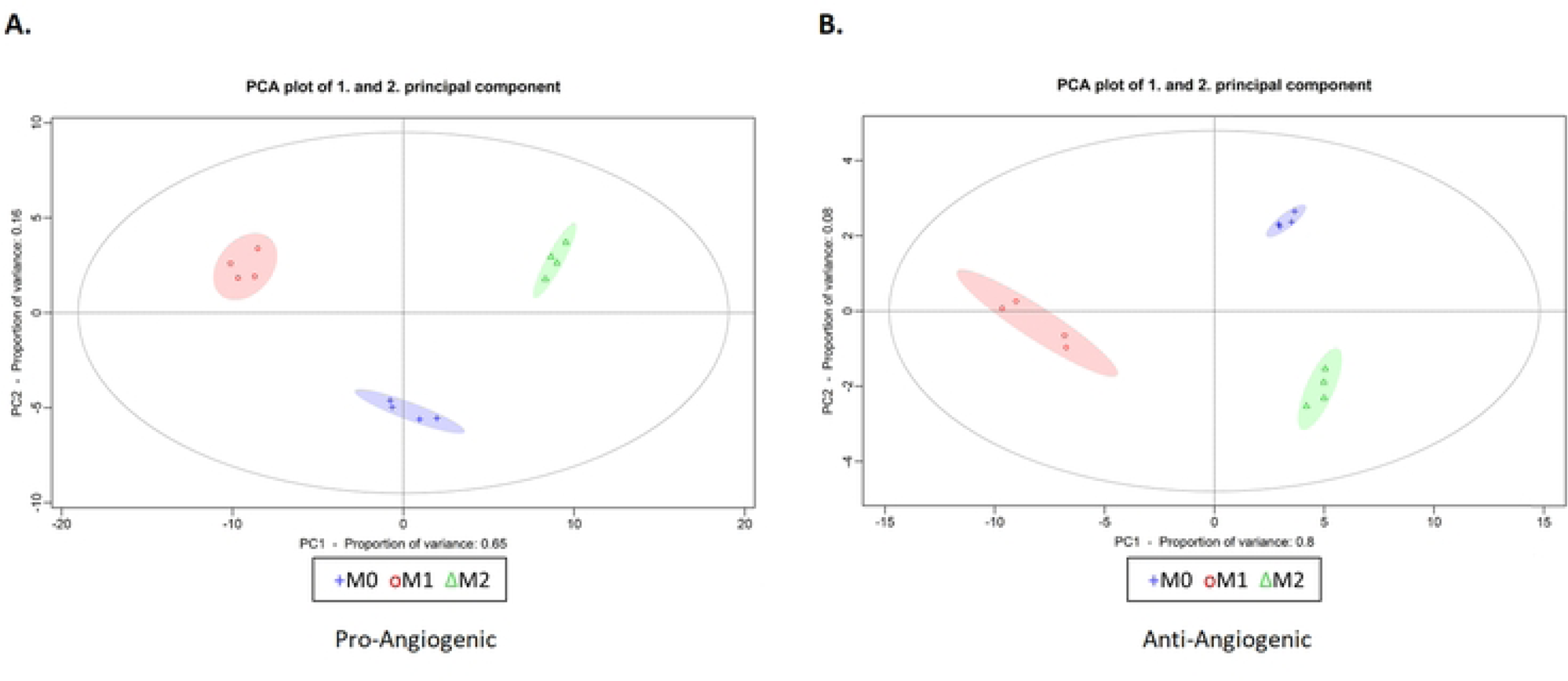
PCA of M0, M1 and M2 macrophages using the expression profiles of pro-angiogenic (A) and anti-angiogenic (B) genes. (A). Both figures show three clusters: blue - M0 cells, red - M1 cells and green - M2 cells.

To enable *in vitro* validation of these *in silico* results, we subsequently selected a set of the most prominently expressed and regulated angiogenesis-related genes out of those used for PCA analysis. Utilizing the PCAGoPromotor package, we ranked these genes based on their individual importance in shaping PC1 and PC2 within each PCA depicted in Fig. 1. This ranking took into account the significance of these genes in both the positive and negative segments of PC1 and PC2, as detailed in Suppl. Table 1. Subsequently, we selected the top three genes from PC1 and the top two genes from PC2 for both the pro-angiogenic and anti-angiogenic gene lists, as outlined in Suppl. Table 1. Notably, we included VEGFA in the list of pro-angiogenic genes, as it serves as a crucial angiogenesis-stimulating factor that exhibited up-regulation in M1 and M2 cells when compared to M0. In annotation, VEGFA is listed as having both pro- and anti-angiogenic activity. Albeit indeed specific VEGF-A isoforms have anti-angiogenic capacity ^24^, we consider VEGF-A production by macrophages primarily as proangiogenic activity.

### *In vitro* validation of angiogenic properties of M1 and M2 differentiation under normo- and hyperglycemic conditions

To evaluate the impact of hyperglycemia on phenotypic characteristics of different macrophage subsets, CD14^+^ monocytes were differentiated under normal (11 mM) or high glucose (25 mM) culture conditions and polarized into M1 (IFN-γ stimulation), or M2 phenotypes (IL-4 or IL-6 stimulation). Bright field microscopy images demonstrated a distinct morphology for IFN-γ-stimulated cultures, characterized by separate colonies of clustered cells that were different from those observed in cultures with M0, or after IL-4- and IL-6 stimulation (Suppl. Fig. 1A). M0 and IL-6-stimulated cultures exhibited a combination of round and spindle-shaped cells, while IL-4-stimulated cultures mostly generated round structures and rarely formed spindles. Flow cytometry analysis showed that under normal glucose condition, IL-6-stimulated macrophages expressed higher levels of CD14, CD16, CD163, and CD206 compared to IFN-γ- and IL-4-induced macrophages, but did not differ significantly from M0 phenotype (Suppl. Fig. 1B). High glucose concentration slightly attenuated the expression levels of these markers in both IL-6-induced and M0 macrophages. IFN-γ-stimulated cells exhibited higher expression of CD64 as compared to the other cell subsets in normal- and to a lesser extent, in high-glucose concentrations. HLA-DR was also expressed at higher levels in IFN-γ- and IL-4-stimulated cells compared to IL-6-induced and M0 macrophages, in both normal and high glucose concentrations.

From these data, it can be concluded that, in agreement with previous findings ^25^, IL-6 stimulation leads to the generation of a distinct subset of macrophages with M2-like phenotype, which exhibit similar morphological and phenotypical features to the M0 subtype. Additionally, IL-4-induced macrophages displayed some overlapping phenotypes with both IFN-γ- and IL-6-induced subsets.

### Both M1- and M2-polarized macrophages express pro- as well as anti-angiogenic factors

To elucidate the relationship between M1/M2 macrophage phenotypes and their pro- / anti-angiogenic properties, as well as the influence of hyperglycemia on these features, we assessed the expression levels of pro- and anti-angiogenic genes in distinct macrophage subsets polarized in normal or high glucose culture conditions using qPCR. These are shown in Fig. 2 and Fig. 3, respectively.

**Figure 2.**
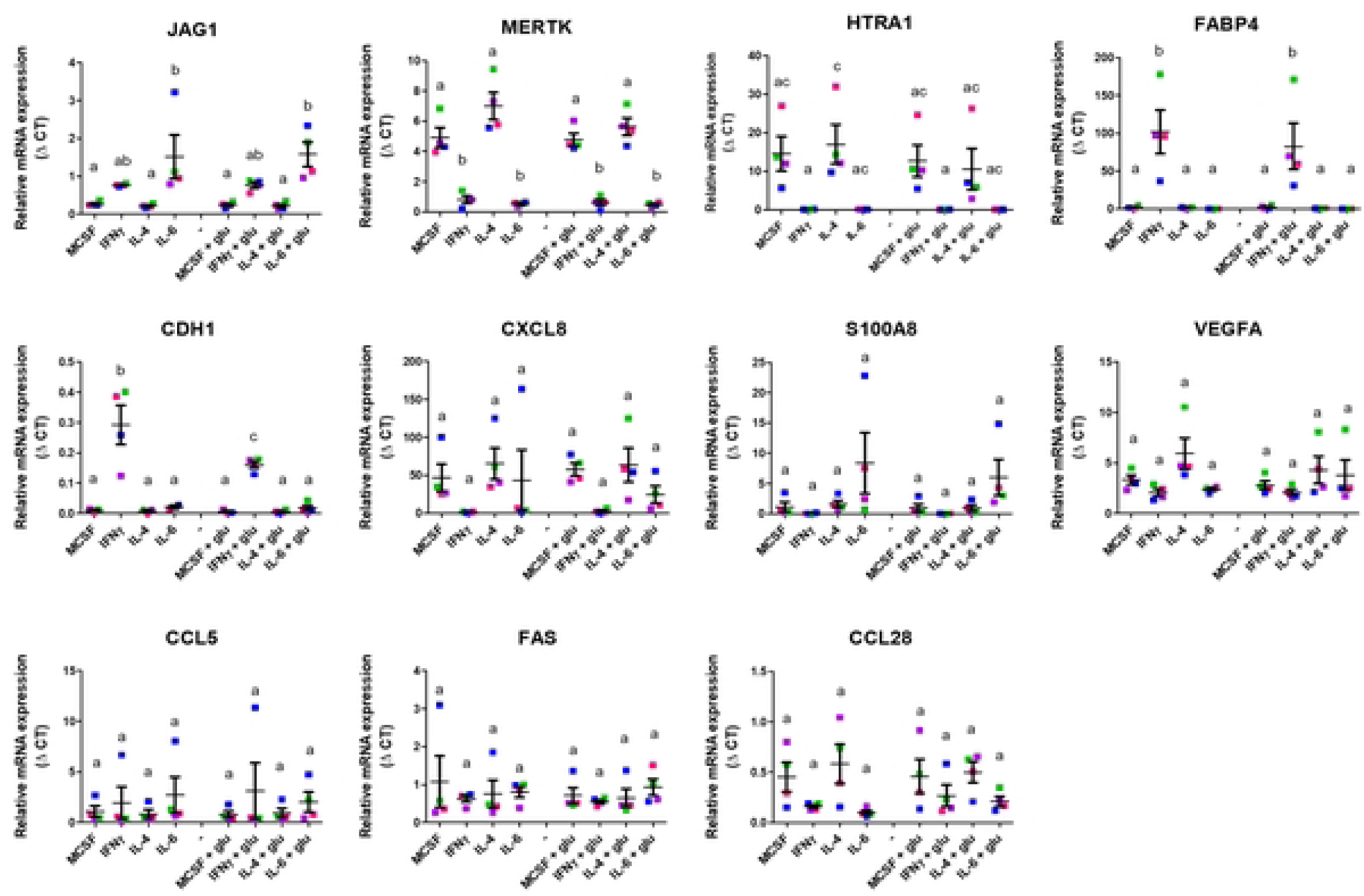
Pro-angiogenic gene expression of M0, M1 and M2 macrophages. Expression of pro-angiogenic genes by different macrophage subsets differentiated in the presence and absence of elevated glucose concentration for 4 days. Expression is depicted as relative mRNA expression. Error bars represent means ± standard error of the mean (SEM). Different letters indicate significant differences. Significance was calculated using one-way ANOVA and Tukey post-hoc test correction for multiple comparisons (*P* < 0.05).

**Figure 3.**
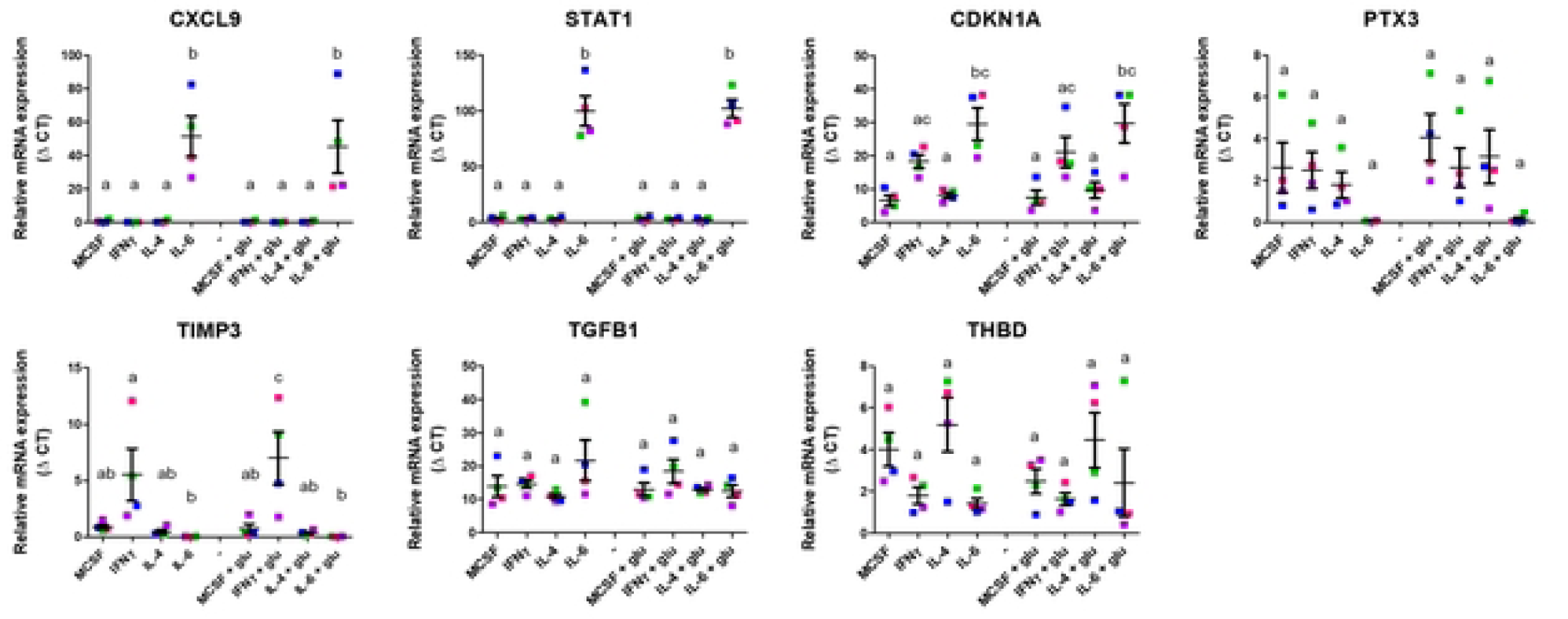
Anti-angiogenic gene expression of M0, M1 and M2 macrophages. Expression of anti-angiogenic genes by different macrophage subsets differentiated in the presence and absence of elevated glucose concentrations for 4 days. Expression is depicted as relative mRNA expression. Error bars represent means ± standard error of the mean (SEM). Different letters indicate significant differences. Significance was calculated using one-way ANOVA and Tukey post-hoc test correction for multiple comparisons (*P* < 0.05).

Analyzing expression of pro-angiogenic genes (Fig. 2), we observed heterogeneous profiles induced by the various conditions. For instance, JAG-1 expression was only significantly elevated in IL-6-induced macrophages compared to M-CSF-stimulated controls. MERTK and HTRA1 expression levels were lower in IFN-γ- and IL-6-induced macrophages as compared to the other subsets. In contrast, IFN-γ stimulation resulted in increased expression of FABP4 and CDH1, and lower expression of CXCL8 (IL-8) and S100A8, although the differences in expression of these latter factors did not reach statistical significance. Furthermore, there were no significant difference in the expression levels of VEGFA, CCL5, FAS, and CCL28 among the different macrophage subtypes, which is at least partly explained by the substantial inter-individual variation.

Among the anti-angiogenic genes expressed by differentially polarized macrophages, similarly diverse profiles were observed, also with variation among individual samples thus limiting statistical power. CXCL9, STAT1, and CDKN1A were higher expressed, and PTX3 expression was lower in IL-6-induced macrophages in comparison with other macrophage subsets, regardless of glucose concentration (Fig. 3). Furthermore, CDKN1A expression was also high in IFN-γ-differentiated macrophages, albeit to a lesser extent than by IL-6. Notably, IFN-γ-induced macrophages exhibited a relatively higher expression of TIMP3. Conversely, no discernable changes in the expression levels of TGFB1 and THBD were observed among the different macrophage subsets. These data represent a complex angiogenic profile that incorporates both pro- and anti-angiogenic characteristics across all M0, M1, and M2 macrophage phenotypes.

To study the integrated results of macrophage phenotype and angiogenesis-related gene expression, we conducted PCA on the combined datasets (Fig. 4). Despite considerable variation, the individual subtypes could be clearly distinguished as clusters based on three distinct components. PC1 primarily discriminated the various polarized subtypes, while PC2 facilitated a separation between macrophages generated under normal and high glucose conditions, with HLA-DR, CDH1, and FAPB4 being the most important contributing factors.

**Figure 4.**
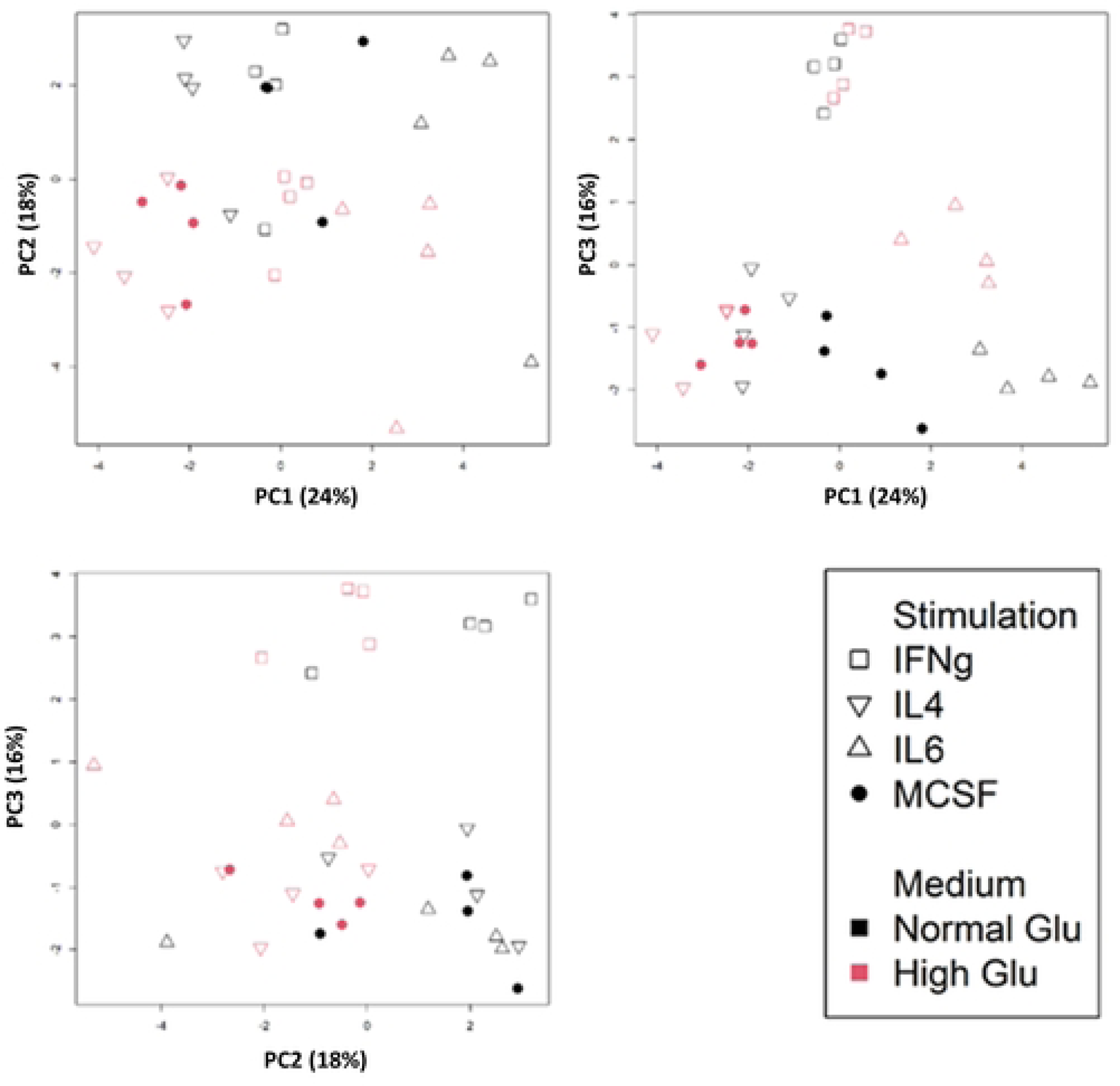
Principle component analysis based on flow cytometry of polarized macrophage phenotypes and qPCR data on expression of angiogenesis-related factors. Three figures show PC1-PC2, PC1-PC3 and PC2-PC3, respectively. PC-values of individual samples are shown. Red color represent high glucose condition.

Remarkably, IL-6-stimulated macrophages exhibited the greatest dissimilarity in PC1 from the macrophages stimulated by IL-4, although both are considered M2-like cells. M0 and IFN-γ-stimulated M1 macrophages take on an intermediate position in this respect. Furthermore, PC3 distinguished clusters that were specific to macrophages induced by IFN-γ stimulation.

### M1- and M2-polarized macrophages exhibit similar, minimal angiogenic capacity in a 3-D in vitro model

To assess the overall angiogenic capacity of the polarized macrophages in a normo- and a hyperglycemic environment, we tested the cells in a three dimensional *in vitro* angiogenesis assay based on co-culture of retinal endothelial cells (REC) and pericytes ^22^ (Fig. 5). We introduced the polarized macrophages in the top medium of the co-culture wells. As positive control for *in vitro* angiogenesis one test condition was always conducted with a mixture of angiogenic growth factors at a high concentration (Hi GF; 200 ng/ml). This concentration of angiogenic growth factors was reduced to 25 ng/ml (Lo GF) in the culture conditions that included the macrophages. In addition, four-day cultured monocyte derived- pro-angiogenic cells (PAC) were used as a positive control as we recently noticed pro-angiogenic capacity of these cells in our *in vitro* 3D retinal angiogenesis model ^26^. Quantitative analysis of the tubules surface area showed that PAC stimulated REC *in vitro* angiogenesis at an equal level to the Hi GF condition and to a greater extent than the Lo GF culture condition (Fig. 5). Although the various macrophage types generally stimulated the formation of marginally increased REC tubule surface areas compared to controls, none of the conditions reached statistical significance compared to the Lo GF cultures.

**Figure 5.**
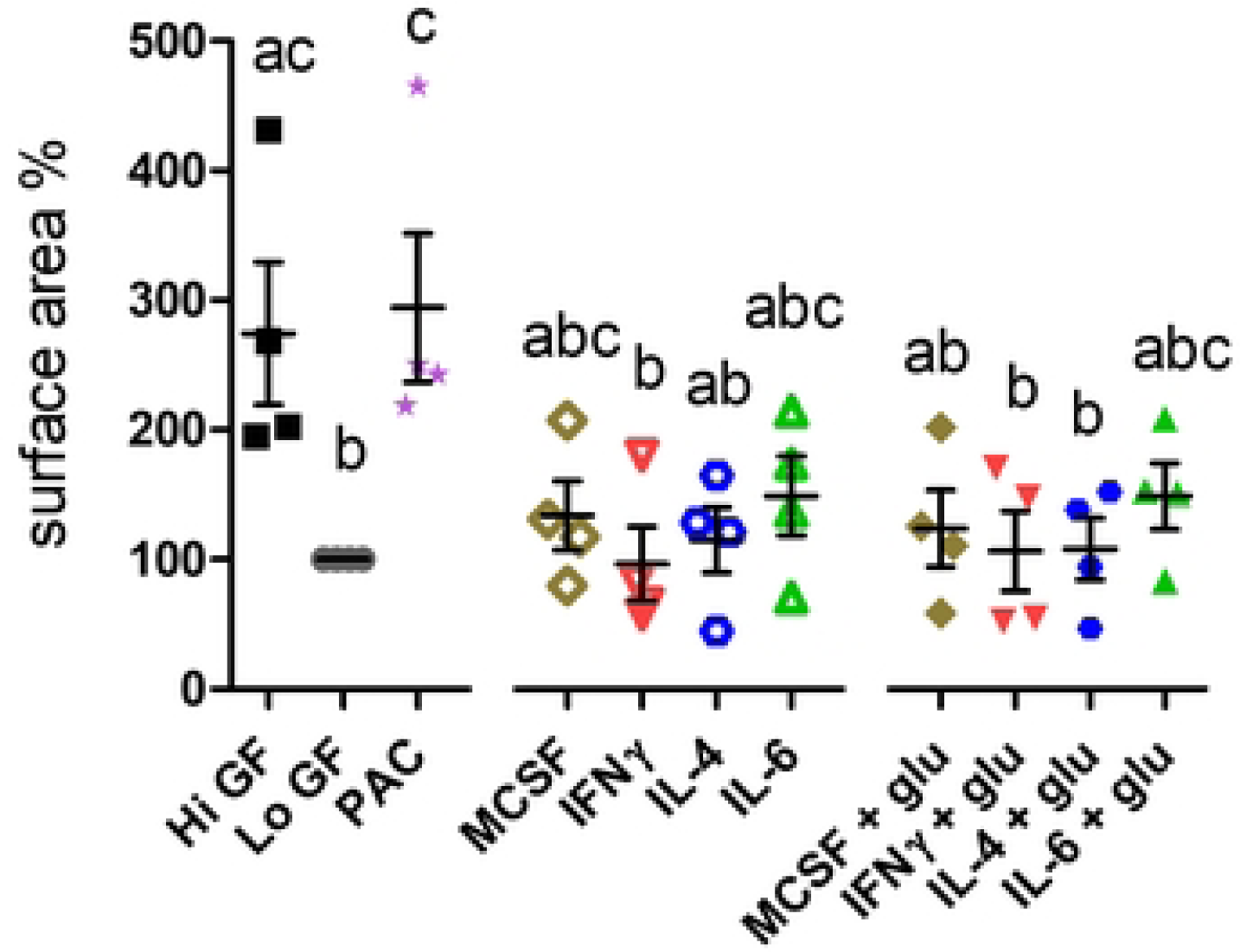
Differentially polarized macrophage subtypes have essentially no pro-angiogenic functionality in a 3D in vitro stimulation assay. In a 4-day culture condition, monocytes were polarized into different macrophage subsets using different stimuli in the presence or absence of glucose. 3D-cocultures of REC and pericytes were stimulated to form tubules using 200 ng/ml (Hi GF) or 25 ng/ml (Lo GF) of angiogenic growth factors in the presence or absence of different macrophage subsets. After 4 days, tubules were imaged and quantified as total surface area percentage, expressed as percentage of total pixels. Error bars represent means ± standard error of the mean (SEM) from four different experiments. Different letters indicate significant differences. Significance was calculated using one-way ANOVA and Tukey post-hoc test correction for multiple comparisons (*P* < 0.05).

## Discussion

In this study, we aimed to investigate in depth the notion that pro-angiogenic activity of macrophages is generally ascribed to M2-polarized macrophages. Furthermore, we examined the impact of hyperglycemia on such macrophage activity, given the prominent angiogenesis-related complications in diabetes. Our findings revealed that both M1- and M2-polarized macrophage subtypes exhibit varying expression levels of pro-angiogenic and anti-angiogenic genes. However, overall angiogenic profiles did not differ significantly between these subsets. Cells generated under hyperglycemic conditions differed from those generated under normoglycemia, based on phenotypic and angiogenic gene expression patterns.

The current constraints of the M1/M2 paradigm in describing macrophage polarization are increasingly becoming evident, as these terminologies delineate opposite ends of a spectrum that are not typically observed *in vivo* ^27^. Transcriptional profiling of human monocyte-derived macrophages has been performed in multiple studies, including by Xue et al., representing a broad spectrum of macrophage activation states extending the conventional M1 vs. M2 polarization model ^8^. It is also suggested that macrophages do not exist as stable subtypes, but are influenced by combinations of factors present in the tissue microenvironment ^28^. Recent studies have investigated how cytokines and pathogen signals impact macrophage phenotypes and challenged the paradigm of distinct subsets based on a limited set of selected ligands in the immune response ^5, 29^. Instead, they suggested that there are pathways that interact to form complex and even mixed phenotypes of macrophages.

Our current study expanded the investigation on macrophage polarization beyond the conventional M1- and M2-activating cytokines, IFN-γ and IL-4, by including IL-6 as an M2-polarizing cytokine ^30^. Our findings revealed that IL-6 stimulation resulted in increased expression of surface markers CD14, CD16, CD163 (scavenger receptor), and CD206 (mannose receptor) compared to M0 macrophages, which is indicative of an M2-like phenotype, in line with earlier literature findings [20,21]. Compared to IL-4- and IFN-γ-stimulated cells ^31^, this profile resembled most closely the M0 subset that also showed low level expression of CD64 and HLA-DR. IL-6 is secreted by various cell types and is implicated in inflammation, response to infection, and wound healing ^32^. It has been reported to increase the expression of M-CSF receptor on monocytes, which is essential for the differentiation of monocytes into macrophages and is known to play a role in the polarization of macrophages towards an M2-like phenotype ^30^.

From the angiogenic gene expression profile in polarized macrophages, we observed an up-regulation of JAG1 (Jagged1), a pro-angiogenic factor and Notch ligand upon IL-6 stimulation^33^. Among the anti-angiogenic genes, STAT1 and its downstream target CXCL9 were up-regulated in macrophages induced by IL-6, both in normal and high glucose conditions. IL-6 activates JAK-STAT signaling upon binding to its receptor, predominantly activating STAT1 and STAT3 ^34–36^. Additionally, we found that IL-6 stimulation increased the expression of cyclin-dependent kinase inhibitor (CDKN1A, also known as p21^Clip1/Waf1^), which regulates cell cycle progression. The up-regulation of CDKN1A in response to IL-6 has been reported by previous studies ^37, 38^.

Based on our data, it appears that IL-6-induced macrophages display a phenotypic profile that shares similarities with M0 and M2-like macrophages, but they tend to manifest a distinct pro-inflammatory phenotype with mixed angiogenic features. The effects of IL-6 can vary depending on the local microenvironment, leading to either pro-inflammatory or anti-inflammatory responses ^39^. Elevated levels of IL-6 have been detected in the microenvironment of various tumors, which are largely driven by macrophages ^25, 39–41^. Together, these studies highlight the complexity of IL-6-induced macrophages and the interaction with the local microenvironment in determining their overall response. Considering both immunophenotyping and angiogenic gene expression of the polarized macrophages, we found that IL-4 stimulation also promotes an M2-like phenotype, which is also similar to M0 phenotype but is distinct from IL-6-induced macrophages as indicated by principal component analysis.

Furthermore, our study revealed a slight inhibitory effect of hyperglycemia on the expression of M2-associated surface markers on macrophages. However, this effect did not translate into changes in the angiogenic functionality of the macrophages. This might be attributed to the short duration of exposure to hyperglycemic condition in our experimental setup, as compared to the chronic hyperglycemia in *in vivo* diabetic conditions, which may lead to epigenetic modifications in macrophages, and altered expression of inflammatory/angiogenic genes. Additionally, it is important to note that our results showed high donor-to-donor variability in terms of gene expression and surface marker characterization. The effect of hyperglycemia on the production of M1 and M2 cytokines in different subpopulations of human primary macrophages has been previously studied ^15^. Moganti et al. have reported that hyperglycemia, in the absence of additional metabolic factors, drives a mixed M1/M2 differentiation profile characterized by production of cytokines that can play a critical role in insulin resistance, diabetes–associated inflammation and vascular complications ^15^. However, to the best of our knowledge, there is limited information available regarding the angiogenic capacity of polarized macrophages under the influence of hyperglycemia.

Our study has certain limitations that should be considered. Firstly, the sample size used in the study was small, which may impact the generalizability of the findings and restricted statistical power. Additionally, studying the effects of hyperglycemia on cellular events using *in vitro* models is challenging due to the relatively short exposure of cultured cells to hyperglycemic conditions. This may not fully capture the complex and dynamic nature of long-term hyperglycemia-mediated changes in macrophage function.

In summary, the findings of this angiogenesis-focused macrophage study support the concept of a spectrum model for macrophage polarization, indicating that the inflammatory and angiogenic status of polarized macrophages is not dichotomous, but rather exists along a continuum. Macrophages exhibit functional plasticity, which is mediated by microenvironmental cues, and can result in diverse phenotypic and functional characteristics, including pro- and anti-angiogenic capacities. However, the underlying mechanisms governing this process are highly complex and necessitate further investigation at multiple levels, including *in vivo* approaches at population as well as single-cell level on gene expression, epigenetics and functionality. Obtaining more comprehensive information in these areas will shed more light on macrophage behavior and facilitate advancements in our understanding of their roles in vascular complications associated with diabetes.

---

**Supplementary Figure 1.**
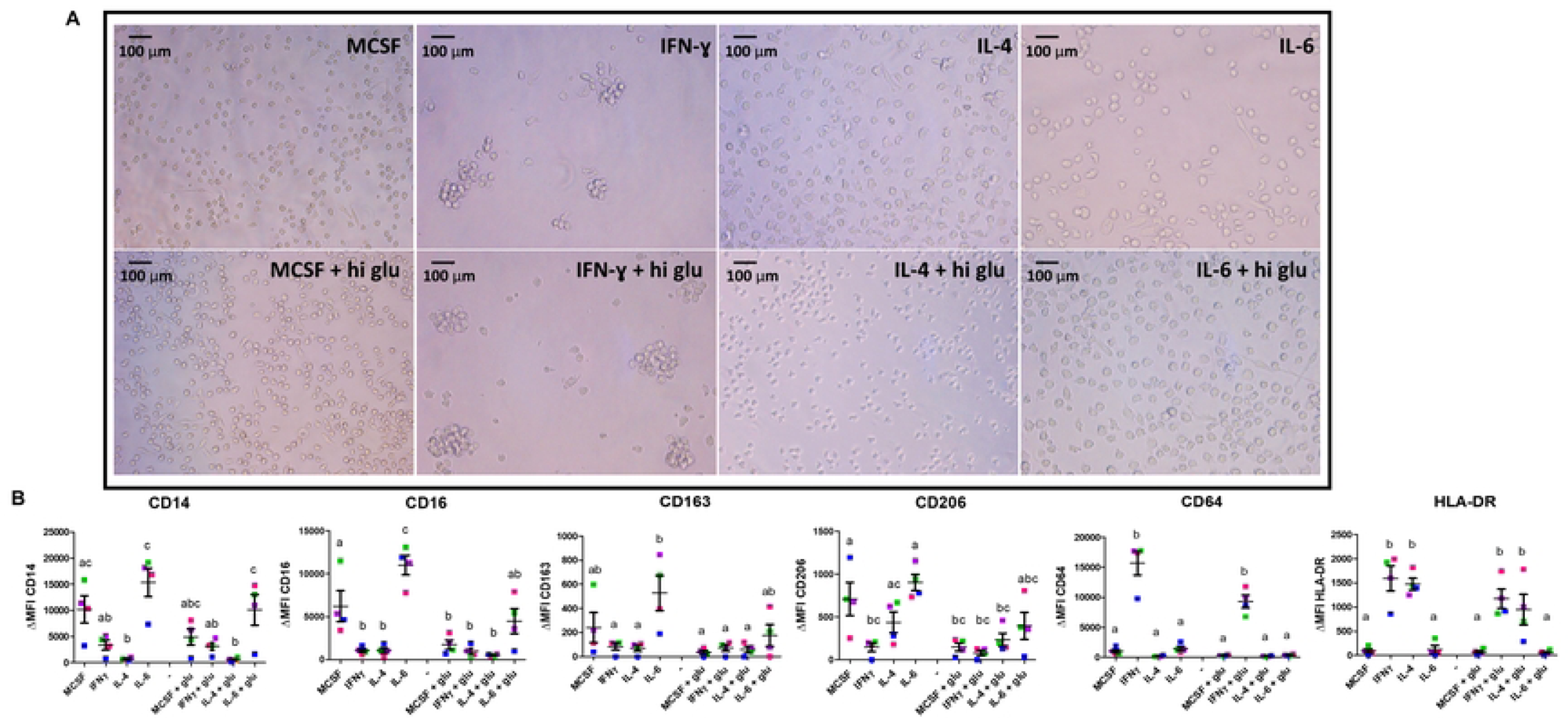
Different immunophenotypes of macrophage subsets; IL-6 drives a distinct M2-like phenotype close to M0. Monocytes were isolated from healthy controls and were stimulated to differentiate into M0, M1 and M2 macrophages under normo- and hyperglycemic conditions. **(A)** Bright field microscopy of different macrophage subsets, 32× magnification. **(B)** Expression of CD14, CD16, CD64, HLA-DR, CD163, and CD206 on differently polarized macrophages. Marker expression was measured by flow cytometry and expressed as median fluorescence intensity (MFI). Values were corrected for auto-fluorescence with the MFI of the backbone control (labeled with CD14 and CD16 only), or unlabeled cells (for CD14 and CD16). Error bars represent means ± standard error of the mean (SEM). Different letters indicate significant differences. Significance was calculated using one-way ANOVA and Tukey post-hoc test correction for multiple comparisons (*P* < 0.05).

**Supplementary Table 1.**
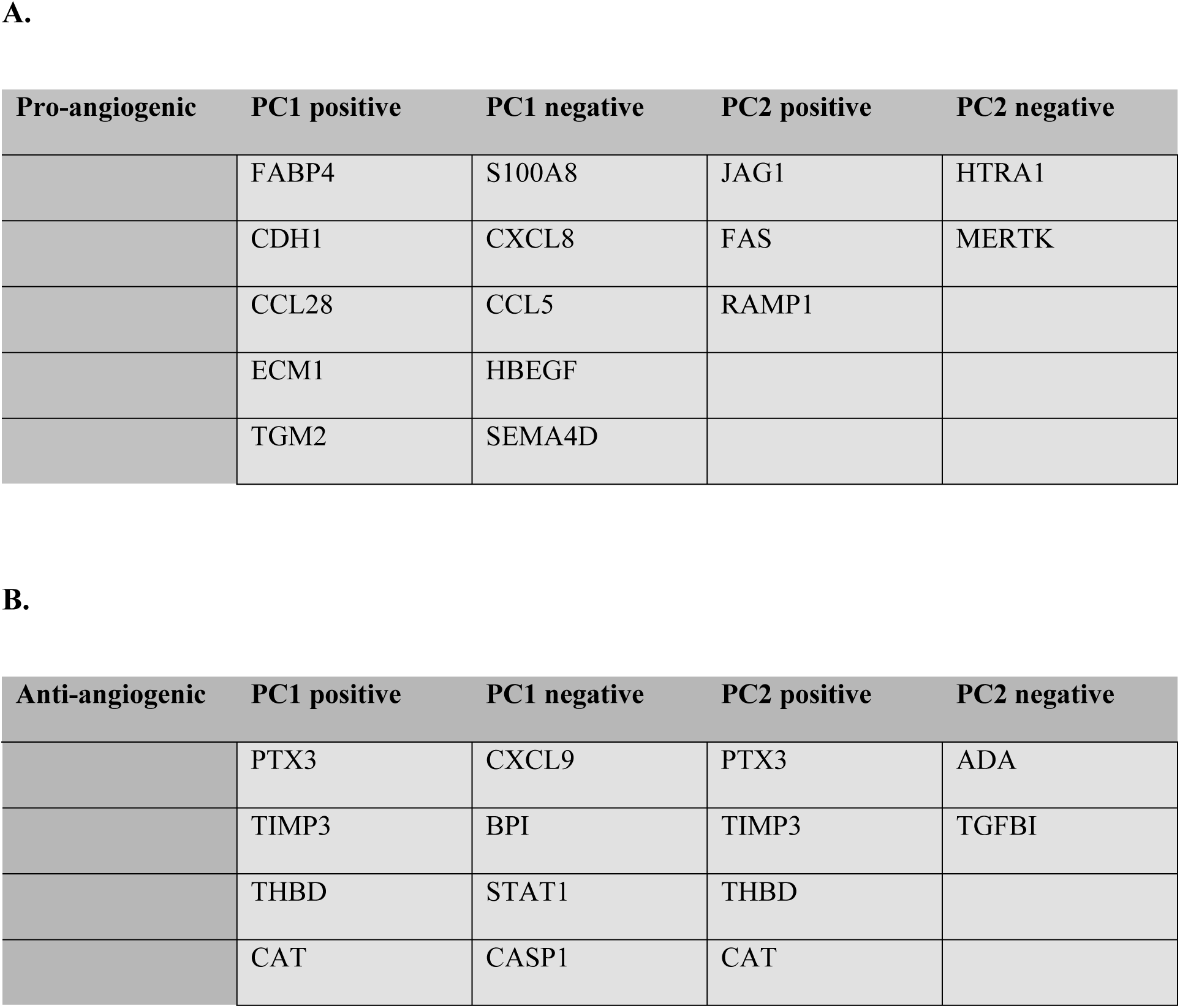
Selected genes for validation experiments. (A) Pro-angiogenic genes (B) Anti-angiogenic genes.

| Pro-angiogenic | PC1 positive | PC1 negative | PC2 positive | PC2 negative |
| --- | --- | --- | --- | --- |
|  | FABP4 | S100A8 | JAG1 | HTRA1 |
|  | CDH1 | CXCL8 | FAS | MERTK |
|  | CCL28 | CCL5 | RAMP1 |  |
|  | ECM1 | HBEGF |  |  |
|  | TGM2 | SEMA4D |  |  |

**B.**
| Anti-angiogenic | PC1 positive | PC1 negative | PC2 positive | PC2 negative |
| --- | --- | --- | --- | --- |
|  | PTX3 | CXCL9 | PTX3 | ADA |
|  | TIMP3 | BPI | TIMP3 | TGFB1 |
|  | THBD | STAT1 | THBD |  |
|  | CAT | CASP1 | CAT |  |

**Supplementary Table 2.**
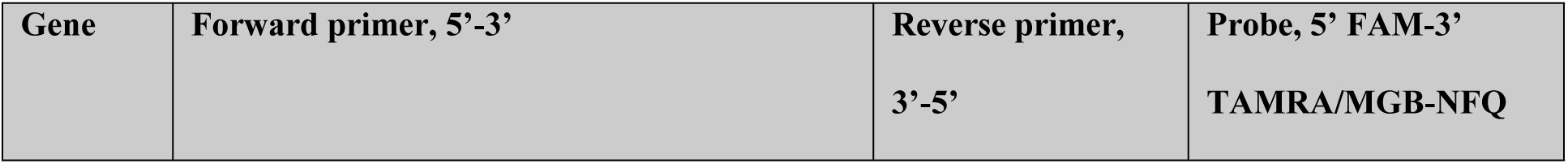
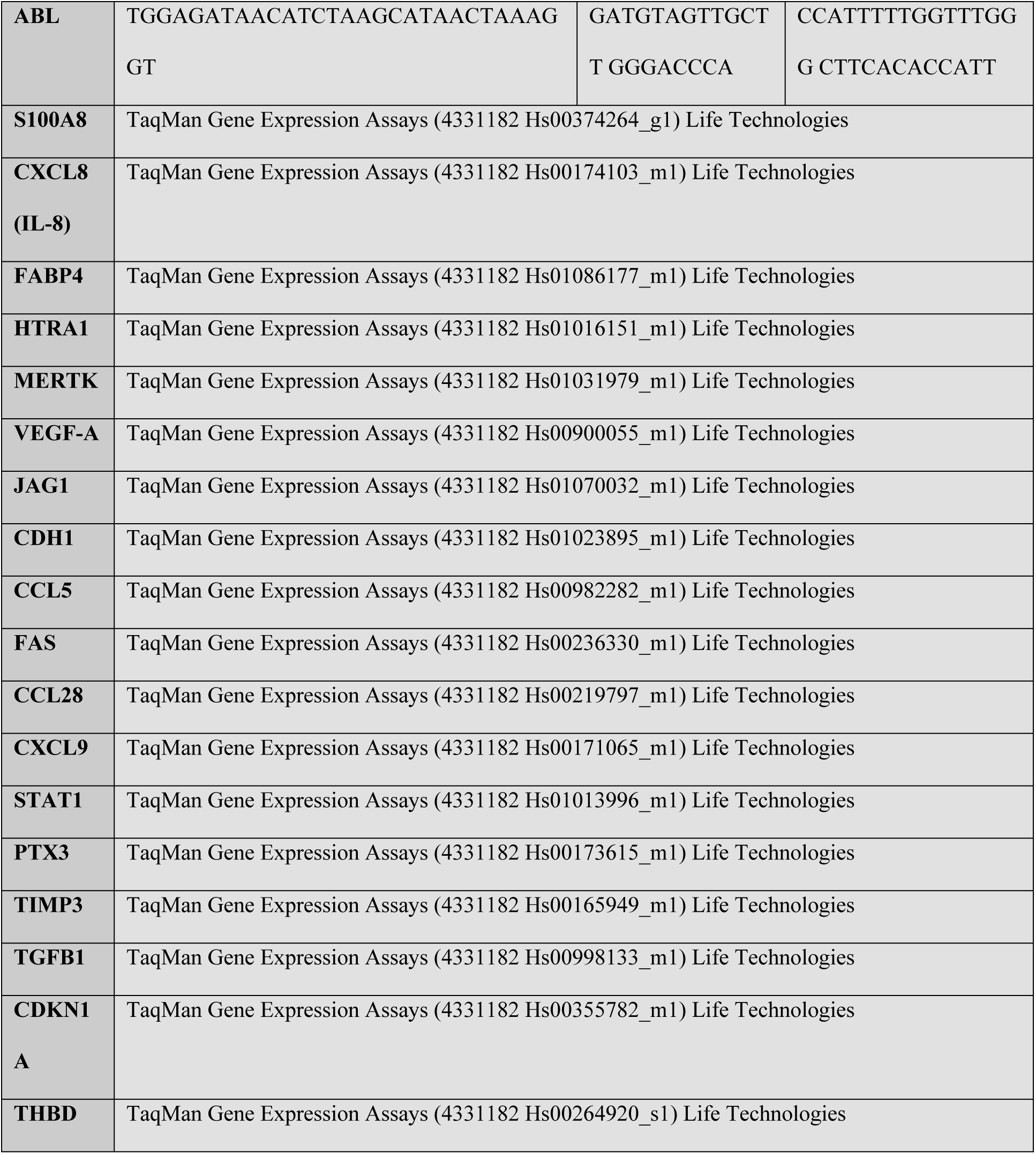
Primer-probe combinations used for qPCR.

**Supplementary Table 3.** Number of DEG in M1 and M2 as compared to M0.

|  | **DEG<sup>b</sup>** | **Up<sup>c</sup>** | **Down<sup>c</sup>** | **Common<sup>d</sup>** |
| --- | --- | --- | --- | --- |
| **M1-NG vs M0-NG<sup>a</sup>** | 1630 | 898 | 732 |  |
| **M1-HG vs M0-NG** | 1753 | 939 | 814 | 1190 |
| **M2-NG vs M0-NG** | 1423 | 663 | 760 |  |
| **M2-HG vs M0-NG** | 1421 | 695 | 726 | 1188 |

*<sup>a</sup>The M0-NG, M1-NG, and M2-NG cells were cultivated using normal concentrations of glucose,* *whereas M1-HG and M2-HG were cultivated with high glucose concentrations.*
*<sup>b</sup>The DEG column indicates the total number of differentially expressed genes observed in the* *comparison.*
*<sup>c</sup>Up and Down columns show the number of up-regulated and down-regulated genes.*
*<sup>d</sup>The first number in the Common column represents the number of DEG that are common to M1-NG* *and M1-HG in comparison to M0-NG, and the second number represents a similar comparison, but for* *M2.*

**Supplementary Table 4.** Pathway analysis of DEG comparing M1 vs M0. (**A)** Regulated pathways in the list of DEG identified in M1-NG vs M0-NG. **(B)** Regulated pathways in the list of DEG identified in M1-HG vs M0-NG. The top 10 regulated pathways and the statistically significant angiogenesis pathways are shown.

| **Order<sup>a</sup>** | **M1-NG vs M0-NG** | **Count<sup>b</sup>** | **Benjamini<sup>c</sup>** |
| --- | --- | --- | --- |
| **1** | GO:0006954~inflammatory response | 80 | 1.69E-13 |
| **2** | GO:0007165~signal transduction | 133 | 0.0019 |
| **3** | GO:0050728~negative regulation of inflammatory response | 21 | 0.0016 |
| **4** | GO:0051056~regulation of small GTPase mediated signal transduction | 28 | 0.0024 |
| **5** | GO:0006915~apoptotic process | 75 | 0.0020 |
| **6** | GO:0006955~immune response | 60 | 0.0020 |
| **7** | GO:0006935~chemotaxis | 26 | 0.0023 |
| **8** | **GO:0001525~angiogenesis** | **38** | **0.0024** |
| **9** | GO:0000082~G1/S transition of mitotic cell cycle | 23 | 0.0028 |
| **10** | GO:0002250~adaptive immune response | 28 | 0.0065 |
| **21** | GO:0045766~positive regulation of angiogenesis | 22 | 0.0262 |

**B.**
| **Order<sup>a</sup>** | **M1-HG vs M0-NG** | **Count<sup>b</sup>** | **Benjamini<sup>c</sup>** |
| --- | --- | --- | --- |
| **1** | GO:0006954~inflammatory response | 72 | 1.68E-09 |
| **2** | **GO:0001525~angiogenesis** | **41** | **4.05E-04** |
| **3** | GO:0006955~immune response | 61 | 0.0014 |
| **4** | GO:0032496~response to lipopolysaccharide | 31 | 0.0045 |
| **5** | GO:0006334~nucleosome assembly | 25 | 0.0058 |
| **6** | GO:0006935~chemotaxis | 25 | 0.0075 |
| **7** | GO:0050728~negative regulation of inflammatory response | 19 | 0.0102 |
| **8** | GO:0007067~mitotic nuclear division | 39 | 0.0099 |
| **9** | GO:0050729~positive regulation of inflammatory response | 18 | 0.0100 |
| **10** | GO:0006915~apoptotic process | 71 | 0.0094 |

<sup>a</sup>The position of the pathway from a smaller *P* value to a larger *P* value.
<sup>b</sup>The number of DEG annotated in the respective pathway.
<sup>c</sup>The corrected *P*-value for multiple comparisons.

**Supplementary Table 5.** Pathway analysis of DEG comparing M2 vs M0. (**A)** Regulated pathways in the list of DEG identified in M2-NG vs M0-NG. **(B)** Regulated pathways in the list of DEG identified in M2-HG vs M0-NG. The top 10 regulated pathways and the statistically significant angiogenesis pathways are shown.

| Order<sup>a</sup> | M2-NG vs M0-NG | Count<sup>b</sup> | Benjamini<sup>c</sup> |
| --- | --- | --- | --- |
| 1 | GO:0006954~inflammatory response | 71 | 1.87E-06 |
| 2 | GO:0051607~defense response to virus | 40 | 7.20E-06 |
| 3 | GO:0045087~innate immune response | 72 | 6.22E-05 |
| 4 | GO:0060337~type I interferon signaling pathway | 21 | 2.06E-04 |
| 5 | GO:0060333~interferon-gamma-mediated signaling pathway | 21 | 0.0011 |
| 6 | GO:0071222~cellular response to lipopolysaccharide | 27 | 0.0017 |
| 7 | GO:0006955~immune response | 65 | 0.0023 |
| 8 | GO:0006919~activation of cysteine-type endopeptidase activity involved in apoptotic process | 22 | 0.0024 |
| 9 | GO:0002576~platelet degranulation | 25 | 0.0022 |
| 10 | GO:0002250~adaptive immune response | 31 | 0.0028 |
| 21 | **GO:0001525~angiogenesis** | **37** | **0.0316** |

B.
| Order<sup>a</sup> | M2-HG vs M0-NG | Count<sup>b</sup> | Benjamini<sup>c</sup> |
| --- | --- | --- | --- |
| 1 | GO:0051607~defense response to virus | 48 | 2.96E-09 |
| 2 | GO:0006954~inflammatory response | 78 | 2.55E-08 |
| 3 | GO:0045087~innate immune response | 79 | 3.50E-06 |
| **4** | GO:0006955~immune response | 74 | 5.98E-05 |
| **5** | GO:0006952~defense response | 22 | 2.48E-04 |
| **6** | GO:0060337~type I interferon signaling pathway | 21 | 4.18E-04 |
| **7** | GO:0060333~interferon-gamma-mediated signaling pathway | 22 | 5.23E-04 |
| **8** | GO:0009615~response to virus | 27 | 0.0027 |
| **9** | GO:0045071~negative regulation of viral genome replication | 15 | 0.0032 |
| **10** | GO:0002250~adaptive immune response | 32 | 0.0040 |
| **24** | **GO:0001525~angiogenesis** | **38** | **0.0475** |

<sup>a</sup>The position of the pathway from a smaller *P* value to a larger *P* value.
<sup>b</sup>The number of DEG annotated in the respective pathway.
<sup>c</sup>The corrected *P*-value for multiple comparisons.

## Notes

### Competing Interest Statement

The authors have declared no competing interest.

